# The energy-saving metabolic switch underlies survival of extremophilic red microalgae in extremely high nickel levels

**DOI:** 10.1101/2024.12.23.630115

**Authors:** Sergio Santaeufemia, Francesca Marchetto, Patrizia Romano, Dorota Adamska, Krzysztof Goryca, Jeffrey Palatini, Joanna Kargul

## Abstract

The red microalga *Cyanidioschyzon merolae* inhabits extreme environments of high temperature (40-56°C), high acidity (pH 0.05–4), and the presence of high concentrations of heavy metals and sulphites that are lethal to most other forms of life. However, information is scarce on the precise adaptation mechanisms of this extremophile to such hostile conditions. Gaining such knowledge is important for understanding the evolution of microorganisms in the early stages of life on Earth characterized by such extreme environments. By analyzing the re-programming of the global transcriptome upon long-term (up to 15 days) exposure of *C. merolae* to extremely high concentrations of nickel (1 and 3 mM), the key adaptive metabolic pathways and associated molecular components were identified. Our work shows that long-term Ni exposure of *C. merolae* leads to the lagged metabolic switch demonstrated by the transcriptional upregulation of the metabolic pathways critical for cell survival. DNA replication, cell cycle, and protein quality control processes were upregulated while downregulation occurred of energetically costly processes including assembly of the photosynthetic apparatus and lipid biosynthesis. This study paves the way for the multi-*omic* studies of the molecular mechanisms of abiotic stress adaptation in phototrophs, as well as future development of the rational approaches for bioremediation of contaminated aquatic environments.

**Importance:** This study provides the first comprehensive analysis of the global transcriptome re-programming in the extremophilic red microalga *Cyanidioschyzon merolae* during its long-term adaptation to heavy metals. We show that the lagged metabolic switch, demonstrated by the transcriptional upregulation of the metabolic pathways critical for cell survival, underlies the long-term Ni adaptation of this model extremophile. The transcriptomic results shed light on how life may have adapted to some of the harshest abiotic stresses on Earth including high temperatures, extreme acidity, and high levels heavy metals that are prohibitive to most other organisms. Additionally, the differentially regulated genes identified in this work provide important clues on the rational development of effective bioremediation strategies of removing heavy metals from the heavily contaminated aquatic environments.

## Introduction

The rapid advancement of the RNA sequencing technologies allows nowadays to gain an in-depth insight into the molecular mechanisms of adaptation to various abiotic and biotic stressors such as exposure to heavy metals. As an example, the transcriptomic studies on the effects of Cd or Ag in a green microalga *Chlamydomonas reinhardtii* showed that genes belonging to metabolic pathways of oxidative stress response and photosynthesis were regulated by these metals (Jamers et al., 2013; Pillai et al., 2014). In a diatom *Phaeodactylum tricornutum*, the analysis of the transcriptome showed that Ni regulated genes involved in metabolic pathways related to the nitrogen cycle, DNA structure remodelling and replication, fatty acids biosynthesis and regulation of the thiol-disulphide redox system (Guo et al., 2022).

Nickel is considered one of the most toxic heavy metals for both eukaryotic and prokaryotic organisms (Das et al., 2019). This heavy metal may be genotoxic, neurotoxic, and even reproductive toxicant. Such effects are dependent on exposure time or the type of contact, whether and how it can enter cells. Specifically, the entry of Ni into the microalgal cells can take place in several forms such as through ion exchange (Liu Ruixia 2002), transient receptor potential-channels (Gómez et al., 2015), or even specific Co/Ni transporters (Blaby-Haas & Merchant 2012). Once inside the cells, it can trigger different responses and effects on the metabolic pathways. In the case of microalgae, the inhibitory effects of Ni constitute mainly the decrease of enzymatic activity, the paralysis of asexual reproduction and the detrimental effects on the structure and function of the main energy ‘factories’ of the cells, the photosynthetic apparatus and oxidative respiratory chain (Monteiro et al., 2012). The main cause of Ni toxicity is its affinity for the -SH groups of enzymes and other proteins, which negatively conditions the growth rate, and the different cellular processes including respiration and photosynthesis.

Considering the high toxicity of Ni and its detrimental effects on cellular metabolism, the knowledge of transcriptome changes caused by Ni exposure is limited, especially in relation to extremophilic microalgae that are naturally exposed to high concentrations of heavy metals such as members of the Cyanidiales order of thermo-acidophilic red microalgae inhabiting aquatic environments of the volcanic origin. A model representative of this fascinating group of extremophilic microorganisms, *Cyanidioschyzon merolae*, was isolated from the volcanic hot springs of Campi Flegrei, Italy (De Luca et al., 1978) and was shown to thrive in moderately high temperatures (up to 57°C), extreme acidic pH (0.05-3) and in the presence of high levels of various heavy metals such as Co, Cd, Hg, Ni, As, Pb, Cr and Fe (Seckbach 2010). Due to its unique evolutionary position at the root of the red algal lineage, this microalga represents a link between prokaryotic cyanobacteria and photosynthetic eukaryotes demonstrated for example through the hybrid characteristics of the photosynthetic apparatus (Miyagishima et al., 2017, Haniewicz et al., 2018). The complete sequencing of its nuclear, mitochondrial, and chloroplast genomes (Matsuzaki et al., 2004), has established *C. merolae* as a model organism for a large number of evolutionary studies and for the dissection of the molecular mechanisms of various fundamental cellular processes including circadian rhythms, lipid and protein homeostasis, cell cycle and division (Kuroiwa 1998; Kuroiwa et al., 1998; Imamura et al., 2015; Miyagishima & Tanaka 2021), and more recently, oxygenic photosynthesis (Busch et al., 2010; Krupnik et al., 2013; Nikolova et al., 2017; Tian et al., 2017; Haniewicz et al., 2018; Antoshvili et al., 2019; Abram et al., 2020; Chang et al., 2020).

In this study, we investigated the transcriptional regulation of various metabolic pathways in *C. merolae* in response to Ni stress. Our work shows that during the long-term adaptation of this extremophile to high Ni concentrations the metabolic switch occurs on day 10 of exposure of the cells to this heavy metal. This switch is underpinned by the transcriptional upregulation of metabolic pathways that are critical for cell survival such as DNA replication, cell cycle, and protein quality control with the concomitant downregulation of energetically costly processes including assembly of the photosynthetic apparatus.

## Results and Discussion

### 1. Dynamic remodelling of C. merolae transcriptome during long-term Ni adaptation

In this study we set out to examine the re-programming of the global transcriptome of *C. merolae* during its long-term adaptation to extremely high concentrations of Ni. The levels of Ni applied in this work (1-10 mM Ni) are up to seven orders of magnitude higher than those present in the natural habitat of this extremophilic microalga (0.003-0.155 mM Ni, Piochi et al., 2019). Figure 1 shows the dynamics of *C. merolae* cell growth under exposure of this alga to 1-10 mM Ni for up to 15 days. As previously showed in Marchetto et al. 2024, the cultures subjected to 1 mM Ni show no significant difference in cell growth compared to untreated control, reaching cell density values of respectively, 2.08·10^9^ and 1.92·10^9^ cells mL^-1^ on day 15. The cell growth was significantly inhibited at 3 and 6 mM Ni from 24 hours onwards, whereby the cell density values of 0.08·10^9^ and 0.02·10^9^ cells mL^-1^, respectively, were reached on day 15. In contrast, exposure of *C. merolae* cells to 3 mM Ni resulted in the initial inhibition of the cell growth in the lag phase (up to day 7, see Fig. 1), followed by the recovery phase between days 10 and 15 (1.05·10^9^ cells mL^-1^ on day 15).

**Figure 1.**
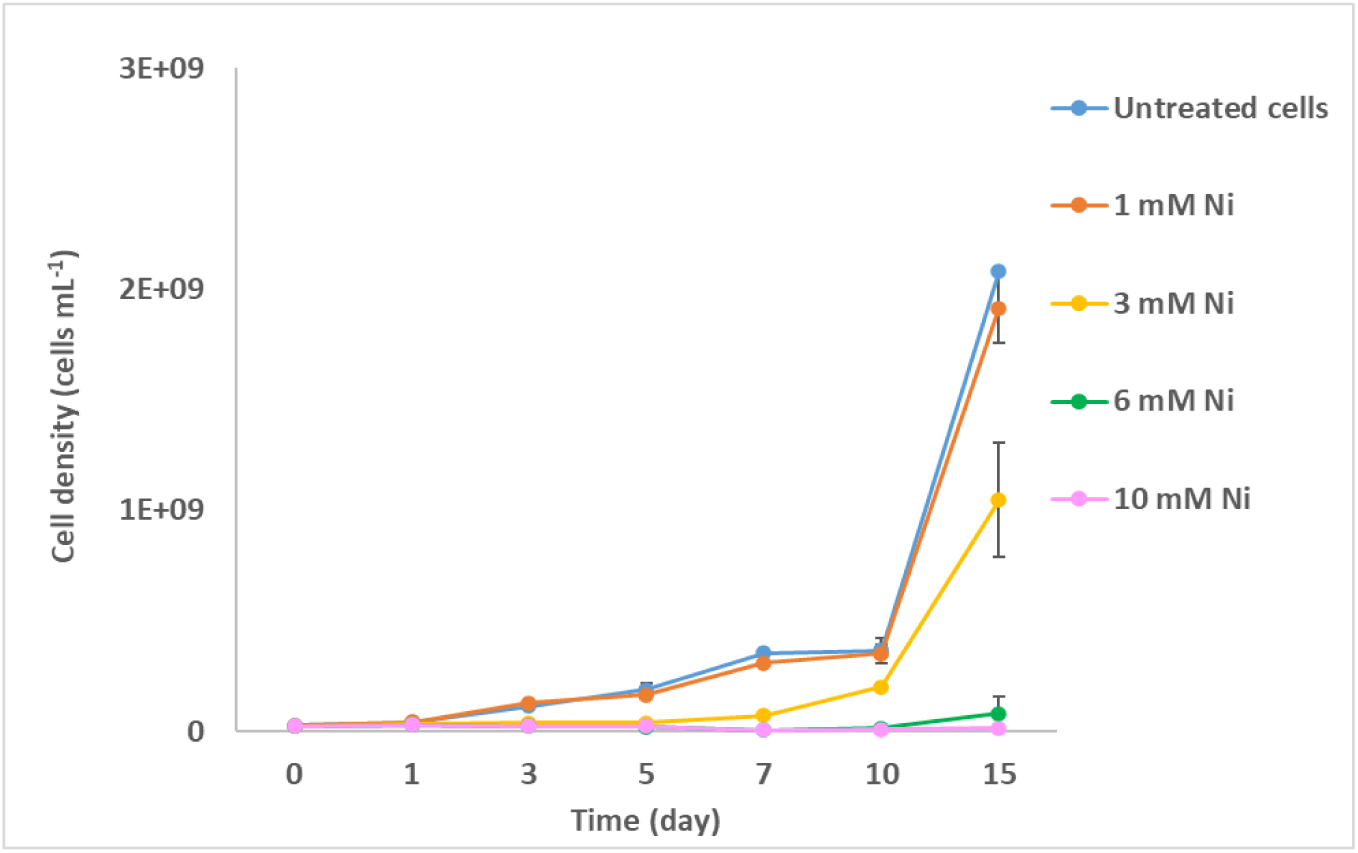
Growth curves of *C. merolae* cell suspensions exposed to various Ni concentrations. Cell density average values (± SD) were determined as described in Materials and Methods for up to 15 days of the Ni exposure for two independent biological replicas (n=2).

In order to gain the first insight into remodeling of metabolic pathways in *C. merolae* during Ni adaptation we performed an in-depth analysis of the global transcriptome changes in response to the exposure of this extremophilic phototroph to Ni levels that yielded cell recovery, i.e. up to 3 mM Ni. This Ni concentration is lethal for most other organisms including the model mesophilic green microalga *Chlamydomonas reinhardtii* (Zheng et al., 2013). To this end, we analyzed the expression levels of a total pool of 4,765 genes at 3 different timepoints (day 5, 10 and 15) during 1 and 3 mM Ni exposure. Of all the considered genes, we identified a total of 252 and 2,888 significantly differentially expressed genes (DEGs; adjusted p-value < 0.05; fold change (FC) > 1 for upregulated or FC between 0 and 1 for downregulated genes) for *C. merolae* cells exposed to 1 and 3 mM Ni, respectively, compared to the untreated control.

The large number of differentially regulated regulated DEGs (Fig. 2A, C) implies the existence of tightly regulated metabolic pathways during the Ni adaptation of the *C. merolae* cells. The over 10-fold higher number of DEGs determined for the cells exposed to 3 mM Ni compared to the 1 mM Ni sample indicates significant remodelling of the *C. merolae* transcriptome during cellular adaptation to the higher concentration of Ni. This phenomenon correlates with the aforementioned cell growth dynamics observed for 3 mM Ni (see Fig. 1).which in turn correlates with the aforementioned cell growth dynamics observed for 3 mM Ni (see Fig. 1) Fig. 2A and Fig. 2C show that the cells treated with 1 mM Ni exhibit the highest number of DEGs on day 5 followed by the gradual decrease at the subsequent timepoints. Interestingly, in the case of 3 mM Ni adaptation, the highest number of DEGs is observed on day 10, which corresponds to the onset of the recovery phase of the cell growth (see Fig. 1).

**Figure 2.**
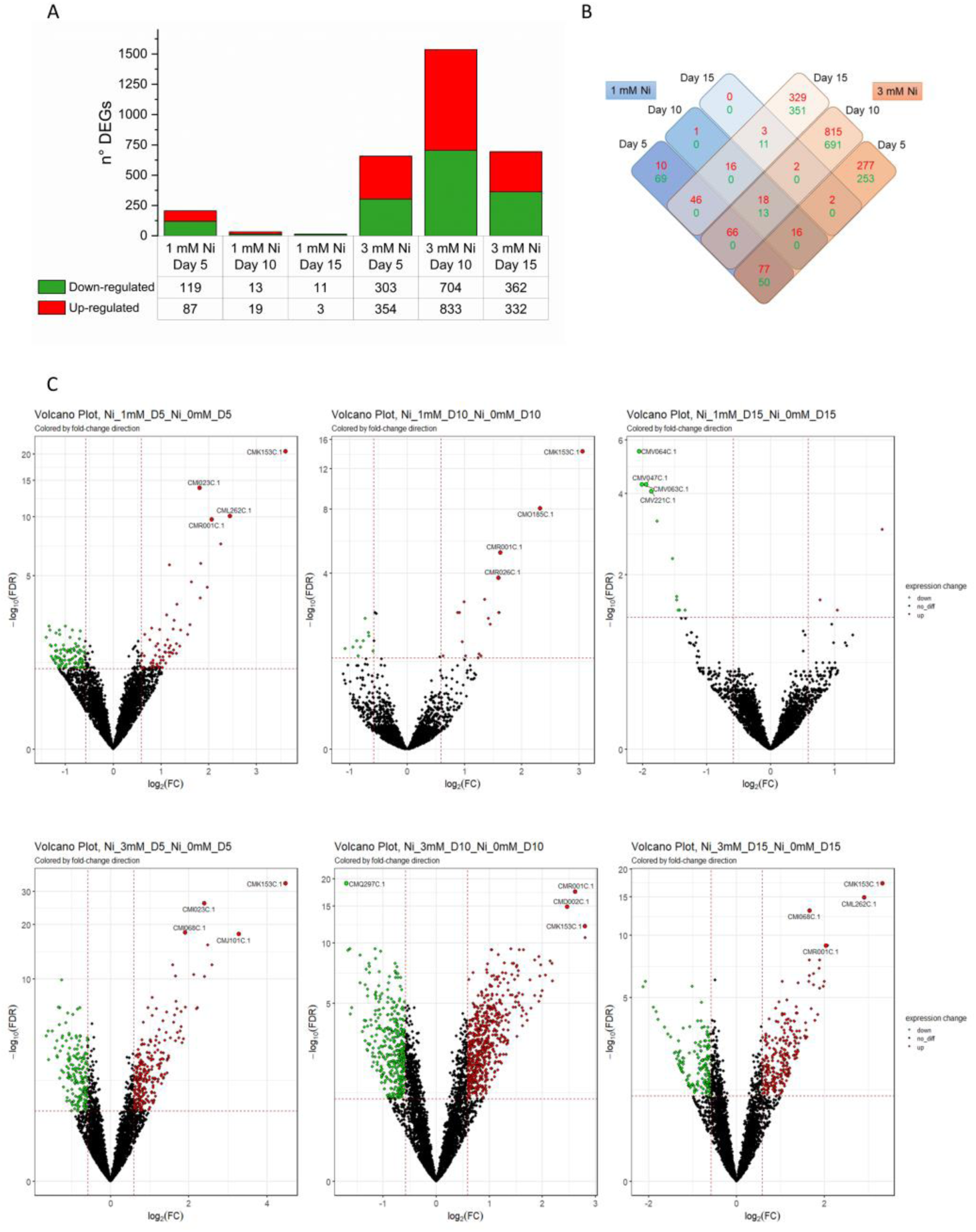
Differentially expressed genes in *C. merolae* cells exposed to 1 and 3 mM Ni. **(A)** Numbers of up- and downregulated genes for the different Ni concentrations and timepoints (5, 10 and 15 days). **(B)** Matrix of differentially expressed genes (DEGs) that were either unique or shared between Ni concentrations tested. Color coding: *red*, upregulated genes; *green*, downregulated genes. **(C)** Volcano plots of DEGs for untreated *C. merolae* cells versus cells exposed to 1 and 3 mM Ni at different timepoints. Plotted are log 10 (FDR) versus log 2 (FC). Data was obtained for three biological replicas (n=3).

The DEGs data shown in Fig. 3 and listed in Tab. S1 implies the existence of the intricate dynamic remodelling of the *C. merolae* transcriptome during long-term exposure to Ni. Differentially regulated DEGs associated with the same metabolic function can be identified at various timepoints of heavy metal adaptation. Below we present a comprehensive analysis of these changes which are related to various types of cellular processes including DNA replication and structural remodeling, cytoskeleton remodelling, oxidative stress responses, intra/extracellular metal transport and sequestration, protein quality control, as well as lipid and energy metabolism.

**Figure 3.**
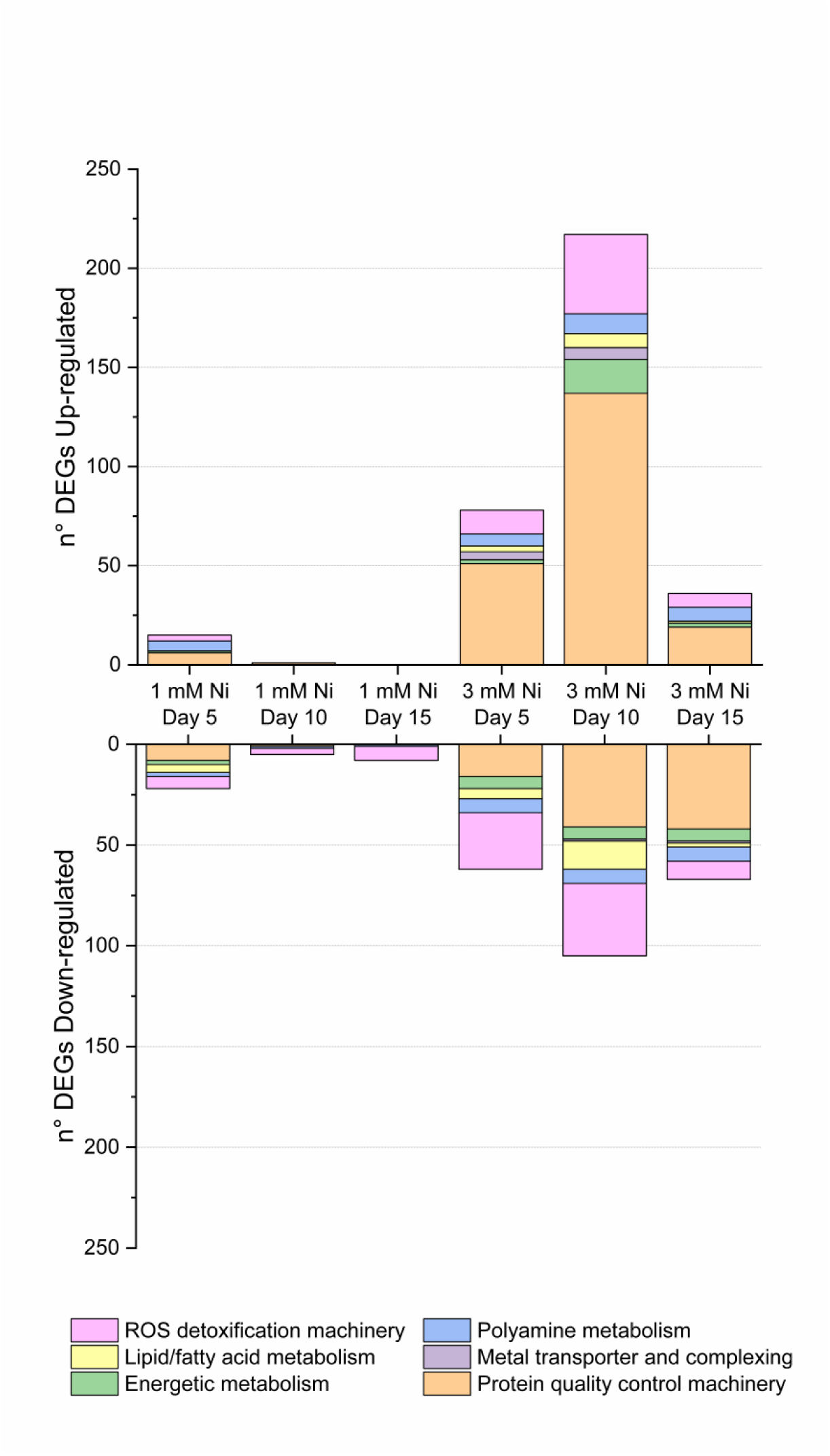
Dynamic regulation of DEGs associated with the specific cellular function during *C. merolae* exposure to 1 and 3 mM Ni. Up- and downregulated DEGs are grouped based on their metabolic function including ROS detoxification pathways, lipid metabolism, energy metabolism, polyamine metabolism, metal transport and complexing, protein quality control and DNA repair.

### 2. DNA replication and structure remodelling during Ni adaptation

As described above, the dynamic global transcriptome remodelling was observed in our study of *C. merolae* cells long-term treated with 1 and 3 mM Ni. In the case of DNA-related transcripts, 1 mM Ni-treated cells exhibited the highest number of DEGs on day 5, whereas for 3 mM Ni-exposed cells, the maximum number of DEGs was on day 10, suggesting different adaptation mechanisms for both treatments (Tab. S2). In particular, on day 10 of 3 mM Ni treatment, the cells expressed the highest number of DEGs related to the regulation of the cell cycle, chromosome segregation, histone modification and DNA repair and replication, whereas on day 15 the highest number of DEGs was related to the DNA replication, showing the different dynamics for the DNA-related transcriptome remodelling during Ni stress adaptation (Fig. 4A). These findings highlight a time-dependent shift in the cellular response to Ni stress, where initial efforts focus on broader genomic stability mechanisms which transition to prioritizing DNA replication processes, underscoring a dynamic adaptation strategy to prolonged metal exposure.

**Figure 4.**
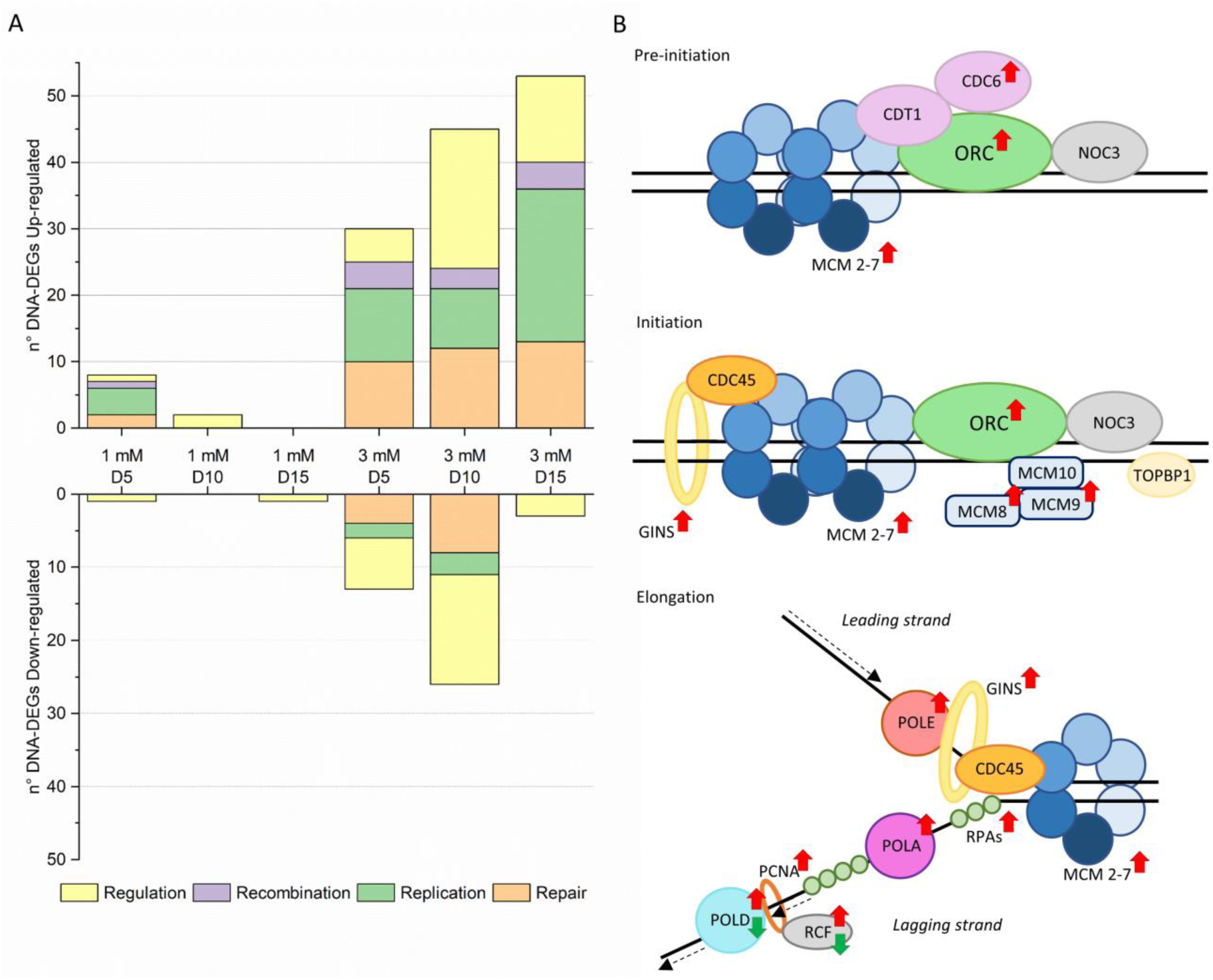
Ni effect on transcriptomic regulation of DNA metabolism. **(A)** DNA related up- and down-DEGs divided by function. The DEGs related to different DNA function are divided in four categories: Repair, Replication, Recombination and Regulation. Repair and Replication group all entries that have a DNA repair or replication function, respectively. Recombination contains DEGs that have a role in the recombination at DNA or chromosomal level, whereas Regulation cluster all entries which function is connected to regulation of different activity, as cell cycle, chromosome segregation, histone modification and DNA repair and replication. In case some DEGs can have multiple function as for example helicase, the entry is reported in all the related possible categories. The figure is based on data reported in Table S2. **(B)** Ni transcriptomic regulation of DNA replication correspond to downregulated once. For specification of the Ni treatment and timepoints see Table S1. Adapted from Shulz et al. (2007).

Heavy metals are known to induce DNA damage resulting in the inhibition of DNA synthesis, increase of mutation frequencies and formation of DNA strand breaks (Nowicka 2022). Moreover, heavy metals in general can evoke damage or aberration at the chromosomal level (Seregin and Kozhevnikova 2006) and inhibition or aberrations of mitosis, cell cycle and differentiation processes (Nowicka 2022). To prevent DNA damage, DNA repair and replication molecular machineries are induced and regulated via chromosome segregation as well as epigenetic changes such as DNA and histone methylation and histone acetylation. Indeed, the level of transcripts related to these processes is significantly upregulated in our study as described in detail below.

During 1 and 3 mM Ni treatment, we observed an upregulation of transcripts linked to the replication pre-initiation complex (pre-RC, Fig. 4B, Tab. S1 and S2), including those involved in replication licensing, like the MCM (minichromosome maintenance) components: MCM1 (CMD025C), MCM4 (CMP365C), MCM5 (CMF173C), MCM6 (CMJ261C), and MCM7 (CMR234C), suggesting a robust cellular effort to prepare for DNA replication during early G1 phase (Tye 1999). This preemptive upregulation likely reflects a stress adaptation mechanism to ensure genomic integrity under Ni exposure. Interestingly, while the pre-RC assembles at various DNA sites, only a subset progresses to DNA synthesis. In our study, the selective upregulation of additional components crucial for DNA synthesis (Shulz et al., 2007), such as MCM8 (CMT087C) and MCM9 (CMO137C), and the entire GINS complex (Psf1 CMR136C, Psf2 CMQ381C, Psf3 CMP212C and Sld5 CMR406C), key players in recruiting the DNA polymerase complex, indicates a precise reinforcement of DNA replication pathways. This implies that cells not only aim to preserve replication readiness but also selectively enhance replication capacity to cope with Ni-induced genomic stress, thereby highlighting a sophisticated regulation of DNA replication dynamics during metal stress adaptation.

In addition to the preemptive replication machinery upregulation, we observed a coordinated upregulation of key components of the DNA replication machinery in response to Ni stress, particularly within the DNA polymerase complexes. Specifically, we noted increased expression of subunits B (CMF127C) and the catalytic subunit (CMI176C) of the DNA polymerase alpha complex (POLA), as well as the catalytic (CMB052C) and B subunits (CMN199C) of the DNA polymerase delta complex (POLD), and subunit B of the DNA polymerase epsilon complex (POLE) (CMH082C). Additionally, replication factor C subunits RFC2 (CMF110C) and PCNA (CMS101C) were significantly upregulated (Fig. 4B). Some of the above-mentioned replication proteins are not only essential for DNA synthesis but also play critical roles in DNA repair, suggesting a dual function in maintaining genomic stability. Notably, we identified upregulation of several DNA repair genes, including those involved in double-strand break repair (e.g., MRE11 CMB035C), mismatch repair (MSH4 CMK199C, CMB035C and MSH6 CMI229C), and DNA excision repair, such as the TFIH family members (CMT290C and CME176C) (Drapkin et al., 1994). The upregulation of the DNA repair genes is also a hallmark of the mammalian systems where increased replication activates MSH/MLH classes of DNA repair genes (Pećina-Šlaus et al., 2020).

The increased expression of these genes suggests that, under Ni-induced stress, cells not only accelerate replication processes but also activate robust DNA repair pathways to counteract potential genomic damage. This dual activation indicates a strategic cellular adaptation to ensure both the continuation of DNA synthesis and the preservation of genomic integrity in a challenging environment.

Another upregulated class of DEGs identified in 3 mM Ni-exposed cells (days 10 and 15) were the components of the SMC (structural maintenance of chromosome) machinery. These proteins form large multiprotein complexes that are essential for chromosomal transmission, condensation and segregation, DNA repair and recombination, as well as epigenetic silencing (Harvey et al., 2002). Notably, genes encoding the SMC1-SMC3 heterodimer (CMI192C-CML027C) and Dcc 1 (CMN029C), which are part of the cohesin complex, were upregulated. This complex plays a critical role in sister-chromatin cohesion during mitosis and meiosis (Strunnikov and Jessberger, 1999), ensuring accurate chromosome segregation. Similarly, components of the condensin complexes, such as SMC2-SMC4 (CMG189C-CME029C),subunit G (CMS422C) and all the subunits of condensin II complex: D3 (CMQ236C), G2 (CMA089C), and H2 (CMI207C), were also upregulated, indicating a reinforcement of chromosomal condensation processes.

Additionally, the upregulation of the SMC5-SMC6 heterodimer (CMH246C-CMA066C), which participates in the ds break DNA repair (Potts et al., 2006), suggests an increased focus on maintaining genomic stability under Ni stress. The correct chromosome alignment and segregation during cell division is fundamental for cell survival and therefore, must be tightly regulated by multiple molecular components. Indeed, in 3 mM-adapted cells, upregulation of genes encoding the kinetochore, a key component of SAC (spindle assembly checkpoint) proteins regulating proper alignment of chromosomes along microtubules during cell division (He et al., 2001) was identified (Tab. S1 and S2). Thus, all four subunits of the kinetochore complex Ndc80 were found to be upregulated in 3 mM Ni-treated cells on day 10: Ndc80p (CMG078C), Nuf20 (CMB012C), Spc24 (CMG186C), and Spc25 (CMI101C) together with Bub1/Mad3 (CMK144C), Mps1 (CMB064C and CMB108C), as well as the spindle and kinetochore-associated protein 1 (SKA1; CMI099C).

Among various downregulated DEGs identified in Ni-treated *C. merolae* cells, the majority appear to cluster around two specific common functions directly related to the regulation of gene expression: histone acetylation and methylation or methyltransferase activity (Tab. S2).This suggests a reduction in chromatin accessibility and gene expression regulation, which could be a response to Ni-induced stress.

Together with the described above upregulation of chromosome condensation and alignment genes, we observed upregulation of the microtubule components of the spindle assembly checkpoint (SAC) in 3 mM Ni-adapted *C. merolae* cells on day 10 and of several microtubule-associated proteins at all-timepoints of 3 mM Ni treatment, indicating a reinforcement of the microtubule network required for mitosis.

Overall, our data implies that cell growth recovery observed from day 10 of 3 mM Ni adaptation of *C. merolae* cells (see Fig. 1) involves at least two levels of upstream transcriptome regulation to safeguard genomic integrity during replication and mitosis: downregulation of the epigenetic modification of histones, in conjunction with upregulation of molecular components involved in DNA repair and replication, as well as chromosome condensation and segregation (see Tab S1 and S2). We propose that these processes constitute the high-level control of the downstream metabolic switch that occurs during 3 mM Ni adaptation of *C. merolae* cells, which in turn provides the basis for the observed cell growth recovery.

### 3. Regulation of oxidative stress responses

Our recent study demonstrated that *C. merolae* cells exposed to 1-10 mM Ni for 15 days display a dynamic amelioration of the reactive oxygen species (ROS) accumulation between day 10 and 15, especially for 3 mM Ni-exposed cells (Marchetto et al., 2024). The extreme environment in which *C. merolae* thrives is rich in several classes of pollutants including heavy metals. Exposure to these stressors can evoke deleterious effects including oxidative stress, indirect formation of ROS and ROS-induced lipid peroxidation (Nowicka 2022). High levels of ROS can ultimately lead to the disintegration of the plasma membrane, deactivation of vital enzymes and apoptosis. The dynamic growth of *C. merolae* in the presence of high (1-3 mM) Ni suggests that during evolution this alga has developed efficient metabolic adaptation to oxidative stress through the activation of a coordinated antioxidant cellular defense response. Our differential expression gene (DEG) analysis revealed key components involved in ROS detoxification during Ni adaptation (Fig. 5, Tab. S1).

**Figure 5.**
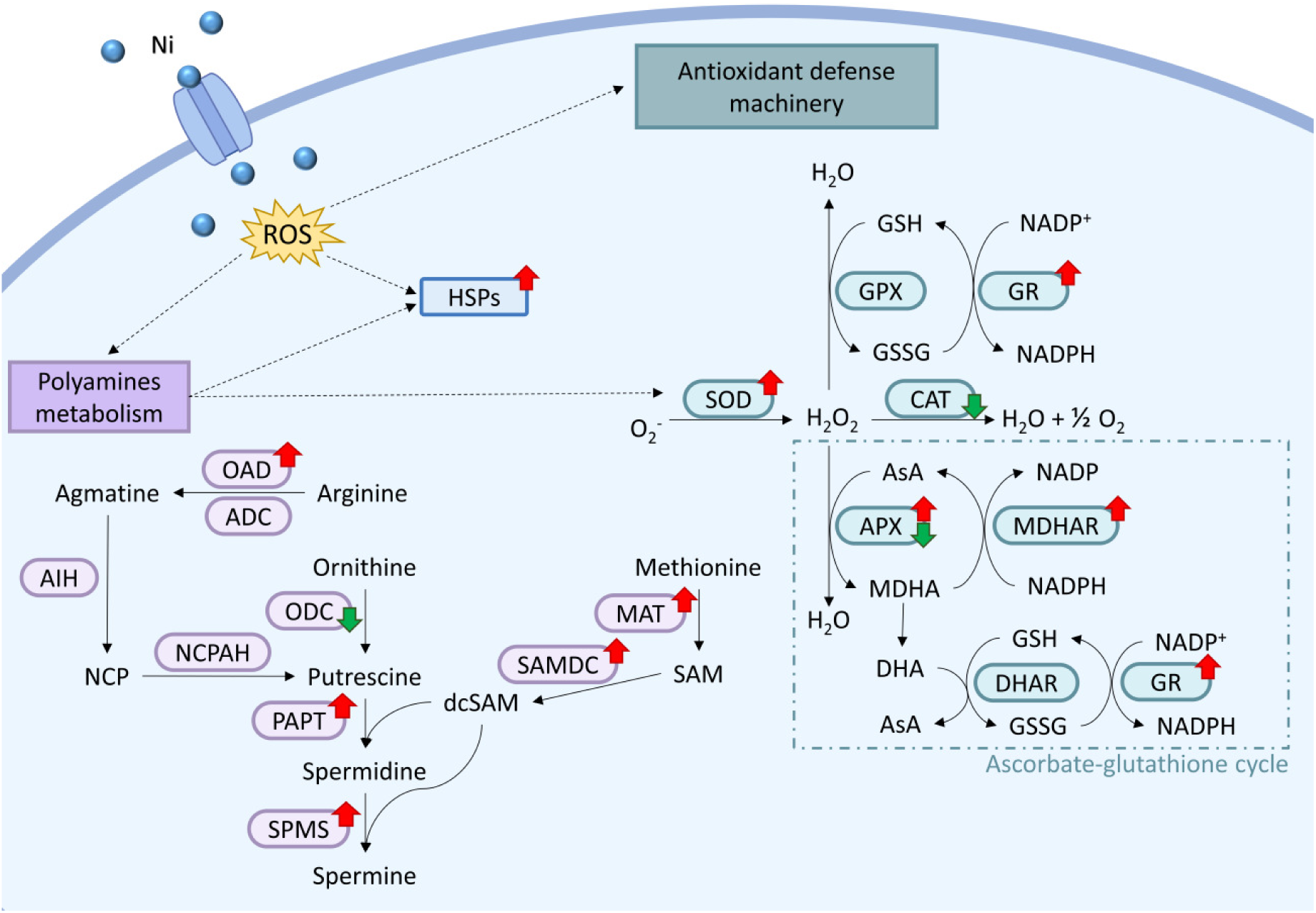
Ni oxidative response machinery in *C. merolae*: antioxidant enzymatic defense and polyamines metabolism. Nickel upon enter in *C. merolae* cells caused an increased in ROS that as signal upregulate the enzymatic antioxidant system and polyamine (PA) metabolisms and heat shock proteins (HSPs). PAs in turn upregulated HSPs and superoxide dismutase (SOD). For detailed timeline of up- and downregulation of displayed enzymes (see Tab. S1). Dotted arrow means that it causes an increase. OAD, ornithine/arginine decarboxylase; ADC, arginine decarboxylase; AIH, agmatine iminohydrolase; NCPAH, N-carbamoyl putrescine amidohydrolase; ODC, ornithine decarboxylase; PAPT, putrescine aminopropyl transferase; SPMS, spermine synthase; MAT, methionine adenosyl transferase; SAMDC, S-adenosylmethionine decarboxylase; GPX, glutathione peroxidase, GR, glutathione reductase; CAT, catalase; APX, aspartate peroxidase; MDHAR, monodehydroascorbate reductase; DHAR, dehydroascorbate reductase; NCP, N-carbamoyl putrescine; dcSAM, decarboxylated S-adenosylmethionine; SAM, S-adenosyl methionine; GSH, glutathione; GSSG, glutathione disulfide; Asa, ascorbate; MDHA, monodehydroascorbate; DHA, dehydroascorbate.

The first line of defense against ROS, Mn superoxide dismutase (MnSOD), is essential converting superoxide radicals into hydrogen peroxide. In *C. merolae*, we observed that a non-specifically localized variant of this enzyme is downregulated on day 10 of 3 mM Ni adaptation, whereas the mitochondrial counterpart is upregulated both on days 5 and 10 (Fig. 5 and Table S1). This observation suggests that transcriptional activation of genes encoding the mitochondrial MnSOD enzyme provides a likely primary response to the increased ROS (e.g. superoxide) levels. This is important in the context of energy preservation metabolic switch proposed in this work in Sections 2 and 6. However, other major ROS-processing enzymes like catalase (CAT), glutathione peroxidase (GPX), Fenton reaction enzymes coupled with the pathway of proline metabolism (Mondol et al., 2022) were either downregulated during 3 mM Ni adaptation or their level of expression remained unchanged compared to the untreated control (Fig. 5, Tab. S1), highlighting the complex fine-tuning of the oxidative stress response during adaptation of *C. merolae* cells to Ni.

Our findings highlight the importance of the ascorbate-glutathione cycle (AsA-GSH) in Ni-induced oxidative stress response. Genes encoding cytosolic ascorbate hydrogen peroxidase (APX), monodehydroascorbate reductase (MDHAR) and glutathione reductase (GR) were upregulated, suggesting that this pathway plays a major role in *C. merolae*’s adaptation to Ni-related oxidative stress (Fig. 5). This pathway seems to be evolutionary conserved as in higher plants, where the overproduction of ROS caused by metal stress was shown to upregulate the enzymes of the AsA-GSH pathway to maintain redox balance (Hasanuzzaman et al., 2019). Interestingly, genes involved in the synthesis of non-enzymatic ROS scavengers such as glutathione (GSH) and phytochelatins (PCs) remain unchanged, although two genes encoding glutathione S-transferase (GST), were upregulated, pointing towards the possible role of these enzymes in the Ni-induced oxidative stress response in *C. merolae*.

During the recovery phase of Ni adaptation, we observed the upregulation of biotin biosynthesis pathways (Tab. S1). Biotin is involved in the formation of thiol peptides such as the above-mentioned GSH and PC, and its upregulation in *C. merolae* mirrors similar findings in another microalga, a mesophilic green alga *C. reinhardtii* under methylmercury stress (Beauvais-Flück et al., 2017), indicating an evolutionary conserved adaptive response to heavy metal-induced oxidative stress. We also observed an intriguing upregulation of mercuric reductase, an enzyme responsible for catalyzes the bio-transformation of Hg^2+^ into Hg^0^ and metacinnabar (β-HgS). These findings along with its activity in another extremophilic red microalga closely related to *C. merolae*, *G. sulphuraria* (Kelly et al., 2007), suggest that mercuric reductase may play a role in managing redox homeostasis also in *C. merolae* upon Ni exposure of the cells, possibly by facilitating the conversion of Ni²⁺ to metallic Ni. This possibility will be explored in future studies.

Another notable phenomenon was the upregulation of polyamines (PAs) biosynthetic genes (Fig. 5, Tab. S1). PAs are known to protect cells from oxidative stress through their ROS scavenging properties. They constitute a group of different molecules that are widespread in all living genera and possess various functions such as nucleic acid protection and regulation of gene expression (Gevrekci, 2017). Moreover, PAs stabilize membranes and metal-binding proteins, potentially reducing intracellular metal accumulation and enhancing stress tolerance (Wen et al., 2010). Therefore, transcriptional upregulation of PAs biosynthesis may result in stabilization of enzymatic antioxidant and metal chelation machineries during mitigation of metal toxicity (Nahar et al., 2016).

Our proposed PA biosynthesis pathway suggests that arginine with a series of downstream reactions, rather than ornithine, might be the precursor for putrescine, the first PA (Fig. 5). Indeed, the downregulation of ornithine decarboxylase (ODC) and the upregulation of enzymes like ornithine carbamoyl transferase suggest the redirection of ornithine through the urea cycle to produce citrulline. Moreover, the upregulation of the OAD enzyme (ornithine/arginine decarboxylase), of which evolutionary studies suggested the main use of arginine as substrate (Lin et al., 2018), should compensate for the downregulation of the ODC enzyme.

In parallel, the upregulation of methionine adenosyltransferase (MAT) and of S-adenosylmethionine decarboxylase (SAMDC) together with the upregulation of PAPT (putrescine aminopropyl transferase) and SPMS (spermine synthase) enzymes indicates a shift toward the synthesis of spermidine (Spd) and spermine (Spm). Both PAs are involved in the amelioration of oxidative stress response especially related to lipid peroxidation (Tajti et al., 2018), inducing the expression of several heat shock proteins (HSPs) (Krüger et al., 2013), increasing the biosynthesis of the photosynthetic pigments (chlorophyll and carotenoids) (Piotrowska-Niczyporuk et al., 2012) and protecting the structural and functional integrity of higher plant thylakoid membranes and photosynthetic apparatus under heavy metal stress (Jiang et al., 2021). Notably, the transcriptional upregulation PAs biosynthetic pathways is consistent with the previously observed increase in photosynthetic pigment content and thylakoid membrane stability in Ni-adapted *C. merolae* cells (Marchetto et al., 2024).

Interestingly, we found that genes related to peroxisomal function were downregulated during Ni adaptation. In the cell, ROS and other oxidative products are formed in the specialized organelles, the peroxisomes (Igamberdiev 2002). These organelles play a role in oxidative stress management and lipid metabolism, yet the majority of peroxisomal genes that play either structural or functional role and key component such as peroxins (PEX) and enzymes involved in catabolic pathway of lipids were all downregulated in 3 mM Ni cells (see data for days 5 and 10 Tab.S1). This observation suggests the induction of a metabolic shift towards energy-saving pathways, redirecting resources from energy-intensive processes like membrane biogenesis and lipid metabolism to essential functions like DNA replication, cell cycle regulation, and ROS detoxification.

In summary, the upregulation of PAs biosynthesis genes and the specific ROS amelioration components identified in this study suggests that these metabolic pathways may be closely related to the previously observed dynamic amelioration of ROS levels in Ni-exposed *C. merolae* cells (Marchetto et al., 2024). On the other hand, downregulation of the genes encoding the peroxisomal proteins and the components of the *β*-oxidation may be related to the above-mentioned metabolic switch into the ‘energy-saving mode’ to limit the energetically costly metabolic processes such as organelle and membrane biogenesis and lipid catabolic pathways and instead, divert the metabolic energy resources towards the pathways that are crucial for cell survival: DNA replication and DNA structural remodelling during cell cycle, regulation of transcription, maintenance of the tight protein quality control and induction of the specific heavy metal-induced ROS detoxification pathways.

### 4. Protein synthesis, folding and degradation

Heavy metals exert adverse effects on multiple cellular components including proteins that can be affected by the direct interactions with the metal or indirectly, through ROS-induced oxidative stress. These effects therefore require the cells to refold the membranes as well as remove and resynthesize the damaged proteins. All these processes are regulated by the vast group of the abovementioned HSP components that ensure accurate protein folding amongst their multiple functions (Park and Seo, 2015). The HSPs play a protective role in various stress responses including heavy metal stress by preventing the aggregation of newly synthesized proteins and priming the damaged/misfolded proteins for degradation (Hasan et al., 2017).

In this work, we observed the upregulation of several HSPs in 3 mM Ni-adapted *C. merolae* cells (Tab. S1), including HSP70 (CMP145C) that was shown to assists in the protein refolding and prevent protein aggregation, leading to increased heavy metal tolerance in higher plants and algae (Hasan et al., 2017, Wang et al., 2004). HSP70 function in concert with a co-chaperone DnaJ also dynamically upregulated in 3 mM Ni-treated *C. merolae* (Tab. S1). Other upregulated HSPs identified in our study in 3 mM Ni adapted cells include nucleotide-exchange factors GrpE (CMT423C and CML088C) and HSP110 (CMS343C), as well as HSP90 (CMA061C and CMQ224C), which has been found to stabilize various intracellular proteins including calmodulin, and cytoskeleton components (Park and Seo, 2015), and CCT protein (chaperonin containing TCP-1 complex), which is required for the production of numerous proteins including actin and tubulin (Brackley and Grantham, 2009). The most broadly upregulated HSP transcript corresponds to a small HSP20 (sHSP20, CMJ101C) which was found to be upregulated for all 3 mM Ni timepoints and for days 5 and 10 for 1 mM Ni treated cells. This HSP acts as co-chaperonin preventing protein aggregation (Park and Seo, 2015) as shown for *Tigriopus japonicus* under different metal stress conditions (Kim et al., 2014).

As heavy metal stress often results in protein damage, it is repeatedly associated not only with degradation of irreversibly misfolded or aggregated proteins but also with enhanced translation to biosynthesize the new properly folded counterparts. Indeed, in 3 mM Ni-adapted cells we observed the massive transcriptional upregulation of the ribosomal proteins including 26 and 38 transcripts coding for different proteins of the small ribosomal subunit 40S and the large ribosomal subunit 60S, respectively (Tab. S1). On the other hand, the transcripts of both small and large ribosomal subunits encoded in the chloroplast genome were downregulated both in 1 and 3 mM Ni-exposed cells at different timepoints (Tab. S1).

Similarly, the differential expression of transcripts coding for several proteasome proteins and for proteins participating in the ubiquitin-meditated proteolysis was observed for Ni-adapted *C. merolae* cells (Tab. S1). Almost all the 22 DEGs of proteasomal subunits for both 20s core particle and 19s regulatory particle, were exclusively upregulated in 3 mM Ni-treated cells, mainly on day 10 of Ni adaptation. On the other hand, the transcriptional regulation of the components of the ubiquitin-proteasome pathway seems to be more complex in the same samples, showing upregulation of the E1-ubiquitin activation enzyme that catalyzes the first step of the process, and either up- or downregulation of the E2 and E3 components involved in the ubiquitin conjugation and ubiquitin ligation steps of the ubiquitin-dependent protein degradation.

In summary, our transcriptomic data for Ni adaptation in *C. merolae* suggests a high demand for chaperonin components required for proper protein folding, and activation of the energy-saving metabolic pathways as demonstrated by the upregulation of the cytosolic ribosomal subunits and downregulation of the chloroplast-localized counterparts. The observed upregulation of proteasomal subunit genes and the up- or downregulation of transcripts encoding the proteins active in the ubiquitin-mediated proteolysis suggest that degradation of damaged proteins may occur in *C. merolae* exposed to Ni mainly in ubiquitin-independent proteolysis. Moreover, the majority of the transcripts related to protein quality control are differentially expressed in 3 mM Ni cells on day 10, when the abovementioned metabolic switch to the metabolic energy-saving mode is likely to occur.

### 5. Lipid metabolism

Oxidative stress, triggered by multiple stress factor such as heavy metals, leads to the peroxidation of lipids that perform a key role in the cell structure as membrane components and in the cell metabolism as an energy reservoir (Nowicka 2022). Changes in membrane properties including fluidity reduction and increase in permeability due to lipid peroxidation lead to a disturbance in the cell organization and function, as observed in our previous study on Ni-exposed *C. merolae* cells (Marchetto et al., 2024).

In this study, we observed transcriptional upregulation of the enzymes involved in fatty acid (FA) elongation in the endoplasmic reticulum (ER) (e.g. KCS, 3-ketoacyl-CoA synthase, CMD118C; KCR, 3-ketoacyl-CoA reductase, CMK172C; and HCD, 3-hydroxyacyl-CoA dehydratase, CMR006C), except for ACCase (acetyl-CoA carboxylase) that catalyzes the first crucial step of conversion of acetyl-CoA into malonyl-CoA. The latter enzyme was downregulated both in the cytosol (CMM188C) and the chloroplast (AccA, CMV056; and AccD, CMV207, respectively). Although, the activity of these enzymes is unknown at this stage and will be the subject of future investigation, it could clarify the different transcriptional levels observed in this study.

The desaturation of fatty acids in phosphatidylcholine, that in *C. merolae* takes place mainly in the ER (Mori et al. 2016), is upregulated in 3 mM Ni-treated cells on day 10 (see data for CMK291C, CMP111C and CMR130C in Tab. S1), along with the transcripts encoding enzymes of phosphatidylinositol biosynthesis (CME109C, CMJ134C and CMM125C, see Tab. S1). On the other hand, the transcripts encoding the enzymes of triacylglycerol (TAG) biosynthesis in the ER (CMR054C, CMN061C and CMQ199C) together with the components of glycolipids and phosphatidylglycerol biosynthesis in the plastid (CMF185C, CMM311, CMR015 and CMV121C) were downregulated (see Tab. S1). In particular, CMV121C component catalyzes the conversion of monogalactosyldiacylglycerol (MGDG) into digalactosyldiacylglycerol (DGDG), the two nonionic lipid constituents of the thylakoid membranes. Another metabolic pathway that is significantly downregulated at the transcriptional level upon Ni adaptation of *C. merolae*, is beta-oxidation of fatty acids (CML080C, CMC137C and CMA042C, see Tab. S1).

In conclusion, the remodeling of lipid metabolic transcriptome in *C. merolae* cells exposed to Ni constitutes an adaptative response to this environmental stress factor: the upregulation of FA elongation and their desaturation may enhance the synthesis of the specific long-chain FA involved in establishing the optimal physico-chemical properties of the membranes such as permeability and fluidity. On the other hand, the transcriptional downregulation of TAG, glycolipids and phosphatidylglycerol biosynthesis can form the adaptive strategy to preserve metabolic energy resources under Ni stress conditions and redirect them towards pathways that are more immediately required for cell survival. In a broader context, changes in lipid metabolism may constitute part of a stress response to ROS that involves the alterations of lipid composition as signaling molecules for the synthesis of antioxidants to counteract the oxidative stress induced by this heavy metal (Sharma et al., 2023).

### 6. Energy metabolism

Microalgae exposed to environmental stress factors, such as heavy metals, can undergo redistribution of energy resources to divert them towards repair, maintenance and defense against the stress itself. The photosynthetic apparatus can be affected by heavy metals at many levels: directly, through the structural and functional destabilization of photosystem II (PSII) and modification in chlorophyll molecules, and indirectly via oxidative stress (Nowicka 2022). A general downregulation of the transcripts encoding the photosynthetic components was observed in *C. merolae* cells exposed to Ni (Fig. 6), including transcripts of photosystem I (PSI), PSII, ATP synthase, ferredoxin, light harvesting subunits Lhcr and cytochromes. The downregulation of photosynthesis genes is a widely used adaptive response in photosynthetic organisms subjected to biotic or abiotic stress (Bilgin et al., 2010; Abdellaoui et al., 2019) to optimize the metabolic energy usage, protecting at the same time the photosynthetic molecular components and avoiding energy imbalance. As a matter of fact, phototrophs can invest the energy resources in immediate adaptive responses necessary for cell survival, without losing photosynthetic capacity thanks to a slow turnover of the photosynthetic proteins (Bilgin et al., 2010). In fact, downregulation of photosynthesis is not always correlated with a loss of function of PSI and PSII, as demonstrated for *C. merolae* cells exposed to 3 mM Ni, where the photosynthetic efficiency of both PSII and PSI complexes remained similar to the untreated control (Marchetto et al., 2024). This may be due to the fact that the production and reassembly of functional photosynthetic proteins are not closely regulated by transcription (Grennan and Ort, 2007).

**Figure 6.**
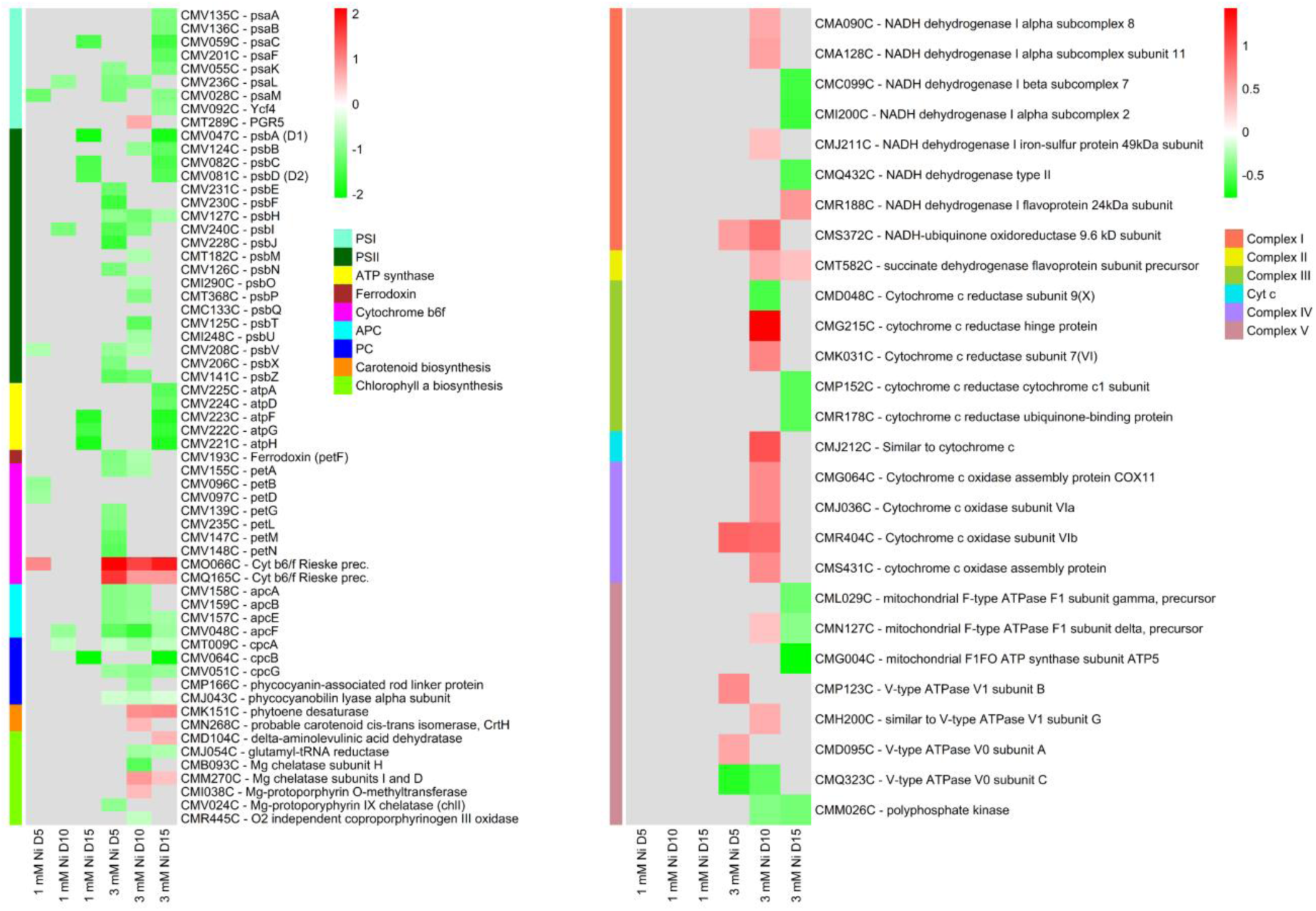
Transcriptomic response of *C. merolae* cells related to proteins involved in energy metabolism. On the left, heatmap of DEGs member of the photosynthetic apparatus or involved in pigments biosynthetic process. On the right, heatmap of DEGs involved in oxidative phosphorylation in the mitochondrion. Data was obtained for three biological replicas (n=3).

Among several downregulated photosynthetic transcripts identified in this study, the notable exception is upregulation of the PGR5 transcript in 3 mM Ni adapted *C. merolae* cells. This protein is involved in cyclic electron flow around PSI and in maintaining redox balance in the photosynthetic electron transport. Other components whose transcripts were upregulated were the two precursors of the iron-sulphur subunit of cytochrome *b*_6_*f*, suggesting a possible adjustment in the composition of the photosynthetic machinery to optimize light energy capture and utilization under the Ni stress.

Previously we showed that Ni exposure affected the total pigment content in *C. merolae*, with the lowest total pigment content measured between day 5 and 10 (Marchetto et al., 2024). Despite the initial reduction of the photosynthetic pigment content, a complete recovery was observed in 3 mM Ni-treated cells, which can be associated with the upregulation of the transcripts encoding enzymes of chlorophyll *a* and carotenoid biosynthetic pathways observed in the present study on day 10 and 15 of Ni exposure and with upregulation of previously discussed PAs.

Our data show that in *C. merolae* upon Ni exposure, the entire glycolysis pathway is transcriptionally upregulated, generally on day 10 of 3 mM Ni treatment, with a few exceptions (Fig. 7) - along with the TCA cycle and part of oxidative phosphorylation (Fig. 6). Upon exposure to various stress factors, microalgae can rapidly generate metabolic energy through glycolysis as a consequence of multiple substrate-level phosphorylation. Besides producing ATP and NADH, both molecules being essential for oxidative phosphorylation, glycolysis serves as a source of precursors for the synthesis of stress-related metabolites and osmoprotectants, supporting the cellular stress response (Ingrisano et al., 2023). The end product of glycolysis is pyruvate, a versatile metabolite that can be directed towards different metabolic pathways. One of them is the tricarboxylic acid (TCA) cycle, in which the carbon atoms of pyruvate are further oxidized for the primary production of high-energy electron carriers, such as NADH and FADH that are subsequently used in the electron transport chain (ETC) of oxidative phosphorylation: the final stage of cell respiration that fuels the ATP production.

**Figure 7.**
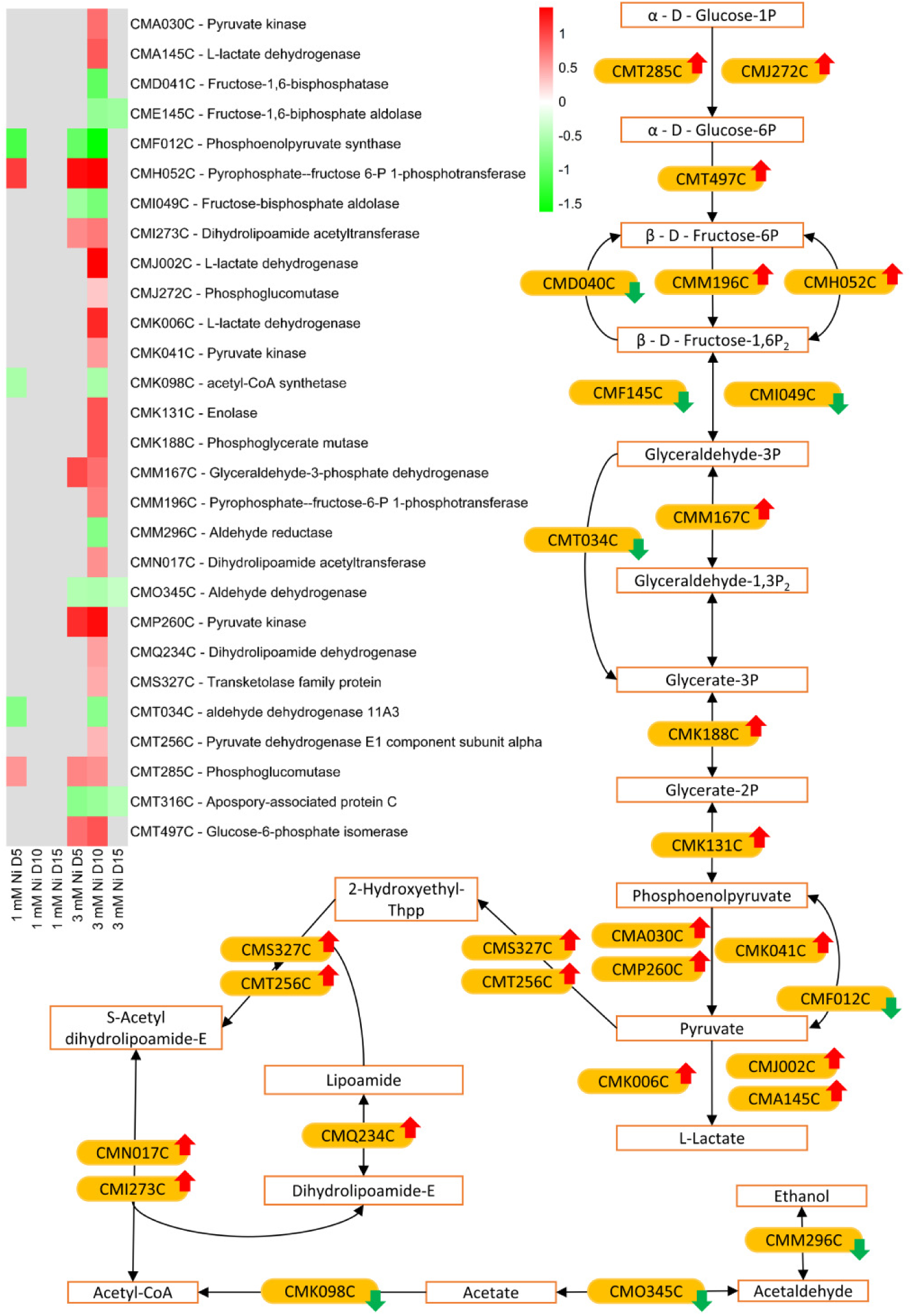
Transcriptomic regulation of glycolysis pathway in *C. merolae* cells exposed to Ni. In the figure based on KEGG glycolysis/gluconeogenesis pathway, the up- and down-regulation of transcripts involved in glycolysis pathway is shown. For detailed time and treatment regulation and for correspondence between accession number and encoded enzyme observed the heatmap on the left and the Tab. S1.

It has been established that upon stress, microalgae may increase mitochondrial respiration as an alternative means of ATP production, in particular if the photosynthetic apparatus is compromised by the stressor. Considering the high photosynthetic efficiency of *C. merolae* cells treated with Ni, the transcriptional upregulation of the alternative energy-generating pathways, particularly on day 10, when a postulated metabolic switch takes place and from which the recovery process begins (Fig 1. and Marchetto et al., 2024), we propose that activation of these processes may be due to additional metabolic energy demand over and above that already produced through photosynthesis.

### 7. Metal transport

In plants and algae several metal transporters were identified that are fundamental for maintaining metal homeostasis; however, in *C. merolae* the characterization of these transporters, their localization and possible function have been determined so far only by bioinformatic analyses (Hanikenne et al., 2005).

Among the different metal transporters present in *C. merolae*, the family of ABC transporters is, without any doubt, the biggest with ubiquitous transporters involved in a several physiological processes. In general, the ABC transporters are mainly downregulated in 3 mM Ni cells on day 10 (see Tab. S1), also considering the ones described in Hanikenne et al. (2005), e.g. CmMRP1 (CMN251C) and CmATM/HMT-1 (CMN105C), which are proposed to transport metal cations from cytosol to vacuole and iron from mitochondrion to cytoplasm, respectively.

Another metal transporter that belongs to a ubiquitous family involved in metal tolerance and homeostasis is the probable manganese transporter CmMTP2 (CMC075C) from CDF family, which is downregulated on day 5 in both 1 and 3 mM Ni -treated cells, and on day 10 only in 3 mM Ni. The CDF, is known to catalyze efflux of transition metal cations from cytoplasm to subcellular compartments or outside the cells.

On the other hand, the transporters CmZIP1 (CMS155C) and CmZIP2 (CMG102C) of the ZIP family, a group of transporters involved in metal homeostasis through the influx of metal cations from outside the cell or from subcellular compartments into the cytoplasm, are upregulated in all the timepoints for 3 mM Ni treated cells, and in the case of CmZIP1 also for 1 mM Ni treatment on day 5.

FTR, NRMAP and IREG1 families consist of groups of metal transporters that mainly regulate iron homeostasis using proton gradient to smoothly transport iron and other divalent cations throughout the cells (Hanikenne et al., 2005). Our analysis shows that this group of transporters seems to be differentially regulated with regards to the specific members. Thus, CmFTR2 (CML004C) and CmNRAMP2 metal transporters (CML262) are significantly upregulated in all timepoints of 3 mM Ni treatment and also in 1 mM Ni-treated cells on day 5. In contrast, CmFTR4 (CMN003C), CmNRAMP1 (CMJ138C) and CmIREG1 (CMG212C) components are downregulated on day 10, and in the case of NRAMP1 also on day 5 in 3 mM Ni-treated cells.

Amongst the DEGs identified in the present study that are differentially regulated during Ni adaptation are Cu transporter CmCOPT1 (CMS307C, upregulated on day 10 in 3 mM Ni), Ca/Mn transporter Ccc1p (CMT466C, downregulated on day 5 and 10 in 3 mM Ni), and Na transporter (CML032C, upregulated on day 5 in 1 mM Ni and in all timepoints of 3 mM Ni). Interestingly, functional protein domain analysis of Na-dependent transporter showed to contain members of the ACR3 family of arsenite permeases which bestow resistance to arsenic by its extrusion from cells (Fu et al., 2009), demonstrating the possible versatility of this group of transporters to various metals. Lastly, the transcript of periplasmic protein p19 (CMG092C) is upregulated on days 5, 10 and 15 in 3 mM Ni exposed cells and on day 5 for 1 mM Ni cells. This protein belongs to the periplasmic metal-binding protein Tp34-type family, which functions together with the importer FTR1, but most importantly is involved in metal sequestration (Brautigam et al., 2012).

The transcriptomic data of metal transporters obtained in the present study does not clarify the previously reported lack of Ni accumulation inside *C. merolae* cells (Marchetto et al., 2024) or whether the transcriptional regulation confers the high metal tolerance, as up- and downregulated DEGs are observed within the same family and with the same predicted cellular localization. Nevertheless, the upregulation of CmFTR2, CmZIP1 and CmZIP2, CmNRAMP2, Na-dependent transporter and the periplasmic protein p19 throughout various timepoints observed in our study points towards the role of these molecular components in maintaining metal homeostasis under heavy metal stress in *C. merolae* cells, perhaps via other metal cation transport systems.

### 8. Other molecular components

Several DEGs coding for trefoil factors proteins were found in this study to be upregulated in both 1 and 3 mM Ni-treated *C. merolae* cells, in particular where the upregulation was apparent for all the timepoints (Fig. 8A). The function of these proteins is still unknown, however, by functional analysis of the protein domains and BLAST analysis, the presence of a P-type trefoil domain, from which the proteins take their name, and a long (circa 500 aa) Kelch-type β-propeller domain suggest the putative protein binding function or a similarity to the signaling Hedgehog proteins. Many of these proteins were found significantly up- or downregulated implying their role in fine tuning of homeostasis during Ni adaptation of *C. merolae* cells.

**Figure 8.**
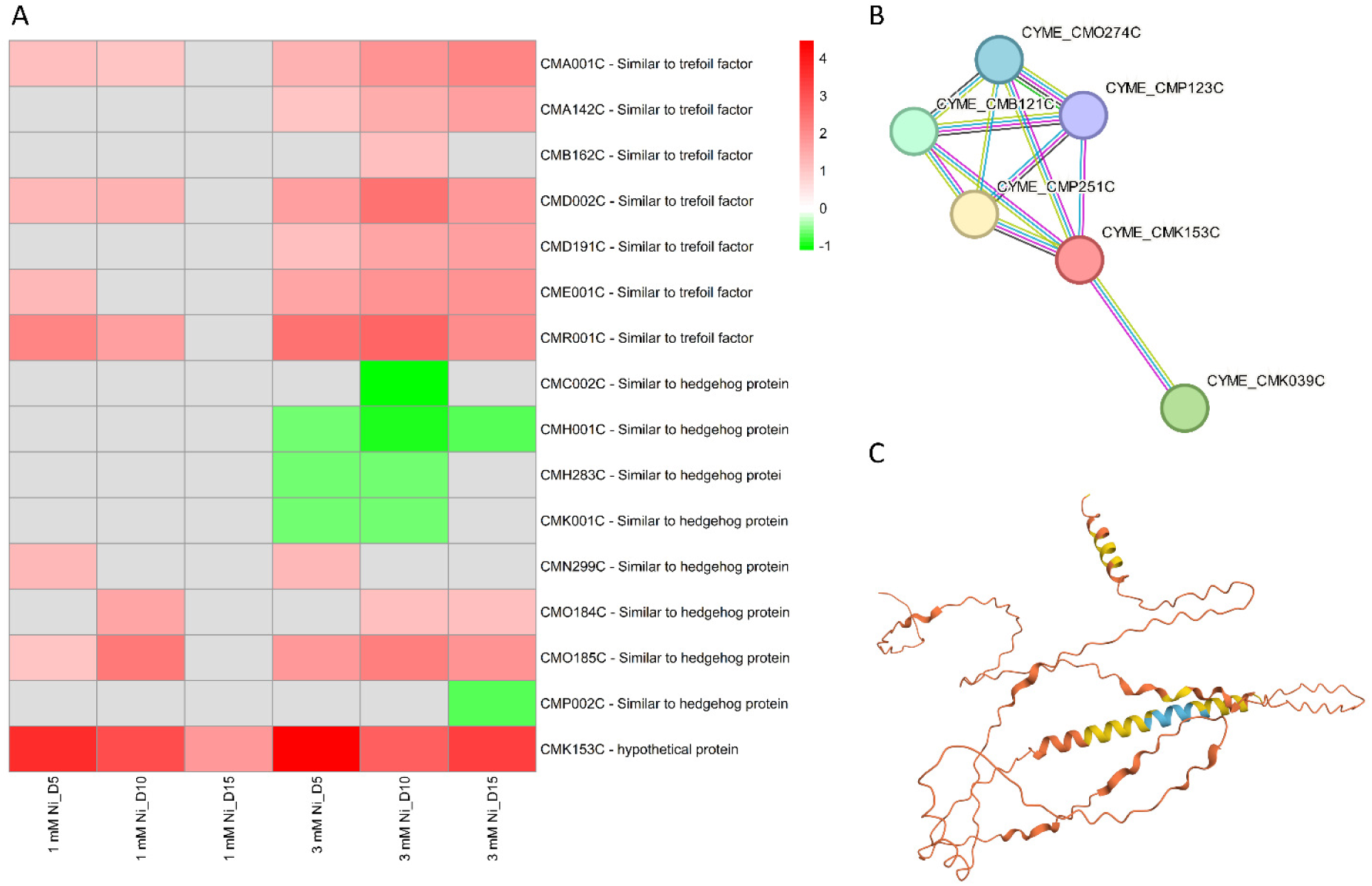
Transcriptomic analysis of DEGs in Ni treated *C. merolae* cells. (A) Heatmap of DEGs mainly involved in signaling pathway. (B) Protein-protein interaction scheme obtained from STRING analysis of the hypothetical protein CMK153C. (C) Structure of the CMK153C protein as predicted by AlphaFold webtool (Jumper et al., 2021). The different color of the protein structure represents the per-residue model confidence score (pLDDT): dark blue - very high (pLDDT > 90), light blue - high (90 > pLDDT > 70), yellow - low (70 > pLDDT > 50) and orange - very low (pLDDT < 50).

An intriguing observation was the transcriptional regulation of the hypothetical protein CMK153C, the only transcript that was upregulated in all the timepoints in both 1 and 3 mM Ni-exposed cells. The bioinformatic analysis gives some clues on the possible function of this protein: the structure prediction, the sequence functional analysis and amino acid alignment suggest the transmembrane nature of this component, with a large non-cytoplasmic extracellular region and two transmembrane *α*-helical regions (Fig. 8C). The hydrophobic region at the N-terminal of CMK153C has some similarities with several transmembrane channel/gate proteins. Moreover, the protein-protein interaction (Fig 8B) shows the connection with V-ATPases (CMO274C, CMP123C), known for their role in pH regulation and proton transport across cellular membrane through ATP hydrolysis; Gtr1/RagA G protein (CMB121C, CMP251C), important for their role in cell growth, nutrient sensing and signaling; and the photoregulatory zinc-finger protein COP1 (CMK039C), which has a role in photoprotection and abiotic stress through signaling pathways (Kim et al., 2022). Considering the possible involvement of CMK153C in light signaling, nutrient sensing, and ion homeostasis, this transmembrane protein may participate in the cellular responses to metal stress possibly with the role in metal transport. Furthermore, the combination of transmembrane localization and interactions with membrane-bound proteins suggests its role in mediating cell membrane signalling pathways. These hypotheses will be verified in future work.

### Conclusions and outlook

This study describes the first global transcriptome analysis of the acido-thermophilc red microalga *C. merolae* in response to extremely high nickel concentrations. It reveals the possible molecular components and pathways underlying the long-term adaptation of this model extremophile to heavy metal toxicity. The DEG analysis revealed that the cells undergo an energy-saving metabolic switch in response to Ni. Metabolic pathways critical for cell survival, such as DNA replication, cell cycle, protein quality control and ROS amelioration, are transcriptionally upregulated, while energetically costly processes, including assembly of the photosynthetic apparatus and lipid biosynthesis, are downregulated. We show that upregulation of molecular components involved in DNA repair and replication, as well as chromosome condensation and segregation are likely to constitute the high-level control of the downstream metabolic pathways differentially regulated in *C. merolae* cells exposed to extremely high Ni levels. This in turn provides the basis for the observed cell growth recovery whereby the optimized metabolic energy expenditure is used for cell survival. Our comprehensive transcriptomic analysis lays the foundation for the future directions in multi-*omic* fundamental studies of stress adaptation in extremophilic microorganisms. Last but not least, the differentially regulated genes identified in this study provide important clues on the possible molecular targets for the development of rational strategies for heavy metal bioremediation in aquatic environments.

## Materials and Methods

### Culture conditions and experimental design

*C. merolae* strain NIES-3377 was obtained from the Microbial Culture Collection at the National Institute for Environmental Studies (Japan (NIES Collection)). The cell culture was cultivated in a Panasonic® Plant Growth Chamber (FL40SS-ENW/37H) using a modified 2x Allen medium (Allen 1959). Cell suspensions (125 mL) were grown at pH 2.5, in continuous white light of 90 µE m^-2^ s^-1^ intensity (400-700 nm), with gentle shaking (110 rpm), at 42°C. These conditions were maintained throughout the experiment. Nickel source used for the stock solution preparation was nickel sulphate hexahydrate (NiSO_4_ 6(H_2_O)) dissolved in MilliQ water.

At the onset of each experiment, the stock *C. merolae* inoculum, NiSO_4_ 6(H_2_O) and an adequate volume of the modified 2x Allen culture medium were mixed to reach the cell suspension optical density (OD) at 750 nm of 0.15. Samples (10 mL) (3 independent biological replicas, n=3) were collected at given timepoints up to 15 days, then were subjected to further analyses.

### Growth measurements

Cell growth was measured as cell density, expressed as cell number (number of cells mL^-1^), and analyzed by counting the number of cells using a Burker Hemocytometer Counting Chamber using an Olympus CX41 light microscope (Olympus, Japan). Based on the cell density, only *C. merolae* control cultures and cultures exposed to 1 and 3 mM NiSO_4_ were selected and considered for transcriptomic analysis.

### Total RNA extraction

Total RNA was isolated from untreated *C. merolae* cells and cells exposed to 1 and 3 mM Ni. A volume of 10 mL of microalgae cell suspension was collected on days 0, 5, 10 and 15 of each experiment. Next, microalgae cell suspensions were centrifuged at 2,200 g for 15 min at 4°C. Then, the samples were frozen and stored at −80°C. RNA extraction was performed using TRIzol reagent (Invitrogen, Carlsbad, CA, USA). RNA concentration was measured using a DeNovix DS-11 spectrophotometer and the purity was assessed by the evaluation of A_260_/A_280_ and A_260_/A_230_ absorbance ratios. In parallel, RNA integrity and quality were assessed with Bioanalyzer (Agilent Technologies, Inc., Santa Clara, United States). Extracted RNA was stored at −80°C until further analysis.

### Construction of RNA-Seq libraries and sequencing

After the RNA quality analysis was performed using Agilent RNA 6000 Pico Kit (Agilent), it was determined that five samples needed additional RNA purification using Kapa Pure Beads (KAPA Biosystems). Enrichment was performed using NEBNext Poly(A) mRNA Magnetic Isolation Module (New England Biolabs). Libraries (mRNA-seq) were constructed using 1 μg of material according to KAPA mRNA Hyperprep (Kapa Biosystems) procedure with KAPA Unique Dual Indexes 96 Plate (KAPA Biosystems). Subsequently, fragmentation was conducted within 5 minutes in 94 °C and were followed by 10 cycles of PCR enrichment. The quality of obtained libraries was analyzed using Agilent TapeStation 2200 with High Sensitivity D1000 ScreenTape (Agilent) and High Sensitivity D1000 Reagents (Agilent). Finally, the quantitative analysis of libraries was measured by qPCR with the Kapa Library Quantification kit (Kapa Biosciences, cat. no. KK48sun), according to the manufacturer’s protocol. Sequencing was performed on Illumina NovaSeq 6000 with NovaSeq 6000 S1 Reagent Kit (200 cycles) (Illumina, cat. no. 20012864), using 100-bp paired-end reads following standard operating procedures and 1% Phix control library addition. As a result, high-quality data was obtained (more than 93 % of data of quality over Phred Score Q37) in a quantity of 4,3-7,3 MR/sample.

### Differentially expressed genes analysis

Raw sequences were trimmed according to quality using Trimmomatic (Bolger et al., 2014) (version 0.39) using default parameters, except MINLEN, which was set to 50. Trimmed sequences were mapped to *C. merolae* reference genome provided by ENSEMBL, (version ASM9120v1) using Hisat2 (Kim et al., 2015) with default parameters. Optical duplicates were removed using MarkDuplicates tool from GATK (McKenna et al. 2010) package (version 4.2.3.0) with default parameters except OPTICAL_DUPLICATE_PIXEL_DISTANCE set to 12000. Reads that failed to map to the reference were extracted using Samtools (Li et al., 2009) and mapped to Silva meta-database of rRNA sequences (Quast et al., 2013) (version 119) with Sortmerna (Kopylova et al., 2012) (version 2.1b) using “–best 1” option. Mapped reads were associated with transcripts from ASM9120v1 database (Zerbino et al., 2018) (Ensembl, version 54) using HTSeq-count (Anders et al., 2015) (version 2.0.1) with default parameters except – stranded set to “reverse”. Differentially expressed genes (DEGs) were selected using DESeq2 package (Love et al., 2014) (version 1.26.1). To consider if one specific gene is differentially expressed, two main criteria were followed: Fold Change (FC) > 1 or < 1, either for an upregulated or downregulated gene, respectively; and adjusted p-value < 0.05. Fold change was corrected using the normal method.

### Gene Ontology and KEGG Pathway Enrichment Analysis

Once the DEGs under nickel exposure of *C. merolae* microalgal cells were determined, a Gene Ontology (GO) and Kyoto Encyclopedia of Genes and Genomes (KEGG) pathway analysis was performed to determine the possible cellular function or metabolic pathway to which each DEG corresponds. The DEGs were then divided into three different categories, such as molecular function (MF), biological process (BP) and cellular component (CC). As in the case of DEGs, Fold Change (FC) > 1 or < 1, either for an upregulated or downregulated gene, respectively; and adjusted p-value < 0.05 was considered significant and used in this analysis. K-means clustering analysis among others were generated and conducted via iDEP 9.6 (integrated Differential Expression and Pathway analysis) webtool (http://bioinformatics.sdstate.edu/idep96/) (Ge et al., 2018).

### Statistical and bioinformatic analysis

For the transcriptomic analysis, three biologically independent replicates of untreated and Ni-treated cells were analysed. Besides, group diagrams were used to display which specific differentially expressed gen is unique or shared between the different nickel treatments and timepoints. On the other hand, Volcano plots and Heatmaps were also generated and created using R as well as principal component analysis (PCA) was carried out to determine the reproducibility between the three biological independent replicates. In the case of up- or downregulated transcripts whose function was unknown or unclear, further bioinformatics analysis were conducted to elucidate their potential roles and functional significance. The bioinformatic resources of InterPro (Bloom et al., 2024), for a comprehensive and integrated analysis of protein sequences and domains, BLAST (McGinnis and Madden, 2024), for the comparison of primary biological sequence, and STRING (Szklarczyk et al., 2023), for the analysis of protein-protein interactions, and AlphaFold (Jumper et al., 2021) were employed.

## Data availability

The data discussed in this publication have been deposited in NCBI’s Gene Expression Omnibus (Edgar et al., 2002) and are accessible through GEO Series accession number GSE284716 (https://www.ncbi.nlm.nih.gov/geo/query/acc.cgi?acc=GSE284716).

## Acknowledgements

This work was supported by the National Science Centre of Poland [MINIATURA 5 grant no. 2021/05/X/NZ2/01516 to S.S. and OPUS 17 grant no. 2019/33/B/NZ3/01870 to J.K.]. The NGS analysis was performed by the Genomics Core Facility at the Centre of New Technologies, University of Warsaw (RRID:SCR_022718), using NovaSeq 6000 platform financed by Polish Ministry of Science and Higher Education (decision no. 6817/IA/SP/2018 of 2018-04-10). We thank Dr. Maurycy Daroch (Peking University, Shenzhen, China) and Prof. Ilaria Corsi (University of Siena, Italy) for fruitful discussions and valuable suggestions on this work.

